# Generative design of synthetic gene circuits for functional and evolutionary properties

**DOI:** 10.1101/2025.09.26.678595

**Authors:** Olivia Gallup, Harrison Steel

## Abstract

In the past decades, a wide suite of design tools for biological systems have been developed, but using these to create biotechnologies that achieve reliable and predictable behaviour remains challenging. Modelling approaches have enabled researchers to traverse the vast search space of genetic circuits more efficiently, while machine learning has proven useful for designing parts and predicting their function or evolutionary properties. Generative algorithms have the potential to leverage these features to design entire genetic circuits from the sequence level, but have only recently begun to be applied to synthetic biological applications. Here, we show that even simple generative models like the conditional variational autoencoder (CVAE) can produce novel genetic circuits that match complex dynamic functions such as signal adaptation. Using *in silico* RNA simulation, we construct a dataset of RNA sequences and convert them to circuits via RNA interaction predictors, allowing us to estimate functional features alongside evolutionary stability and interpret model-learned features. Our model generates diverse distributions of circuits that match their target adaptation specification well, even when limited to small training data sets. Structures in the embedding space correspond to motifs previously identified as crucial for adaptation and reflect the design rules for adaptable circuits. Framing adaptation as a single design objective outperforms other input representations, reflecting the importance of choosing the correct data encoding for generating genetic circuits. Finally, we show that functional and evolutionary properties can be prompted simultaneously, providing a proof-of-concept for the combined design of phenotype and evotype.

## 2 Introduction

Biological systems must constantly adjust to changes in their environment and return to homeostasis after perturbations. Adapting to an external signal involves returning a system back to its original state before it was perturbed; this functionality is widely used in natural systems where precise sensing or control is required [1–4], and can be realised by a surprisingly small number of interacting components in an organism [5, 6]. In synthetic biology, designing genetic circuits that can adapt to disturbances is crucial for stable functioning, as biodesigns are applied in different applications or environments [7, 8].

However, biological circuits can fail for many reasons [6, 9, 10], such as unforeseen host interactions, parameter-fragile designs, or mutations. As an example, designing a circuit that adapts to perturbations can be achieved with motifs such as incoherent feed-forward loops or negative feedback with a buffer node [5, 11–14], yet functionality is often disrupted by host context and *in general* constrained by practical requirements (such as non-repetitive sequences, low crosstalk, assembly compatibility, and synthesis ease). Design optimisation can encode such requirements in objective (cost) functions [15–17], but some such properties entail significant computational hurdles, such as characterising mutation effects and the surrounding evolutionary landscape. Meanwhile, changing the target specification (e.g., desired adaptation level) often forces a fresh search.

Currently, designing evolutionary properties is most important in directed evolution, where starting variants must be functional yet still evolvable. In synthetic biology, the aim is typically mutation resistance – coupling function to an essential process [18] or imposing growth costs on burden-reducing mutations [19]. Nevertheless, the importance of tuning a circuit’s evolutionary “evotype” alongside its physical “phenotype” is becoming increasingly recognised [20]. In nature, distinct functions that are separated by small evolutionary distances allow populations to adapt quickly to new environments. A study on the LacI repressor in *E. coli* found that a few mutations could transform its function entirely, yielding inverse dose response or a band-pass behaviours, all only a few amino-acid steps away [21]. In this way, the evolutionary landscape around a genetic circuit can hold a plethora of hidden functional possibilities.

To make building genetic circuits more predictable, researchers developed a variety of mathematical modelling approaches. Algorithms for exploring possible architectures often rely on parameter-level approaches, such as exploring biological part libraries [22, 23], optimising directly from a starting set using gradient descent [24], or probabilistic approaches [25, 26]. Alongside optimisation, other approaches focus on building predictors, such as physics-based simulators that determine RNA structure and interactions [27, 28]. Some tools utilise nucleotide or amino acid sequences directly [28–30] and have become standard in synthetic biology [31, 32] [33, 34].

Recently, machine learning (ML) predictors have become widely used for their versatility and accuracy. In biology, applications include tuning expression levels [35–38], building functional switches [39], and linking composition to function [40, 41]. These models can be integrated into existing tools for building complex circuits, such as logic gates, from libraries of characterised biological parts [42–47]. Beyond prediction, generative ML models have been used to propose entirely new or modified parts, such as designing DNA and RNA sequences to boost transcription [36, 48] and translation [49, 50], creating diverse protein libraries [51–53], and elucidating metabolic function [54] and mutation effects [35, 55]. Hybrids of these optimisation, physics-based, and generative approaches could markedly improve the reliability of biodesign and be useful for downstream applications [56].

So far, generative models have been deployed in synthetic biology to design individual parts [48, 57], with the aspiration of designing entire circuits [40, 58]. Optimising biological parts (riboswitches, promoters, or repressor proteins) typically requires a model to recognise sequence features that determine the binding interaction. However, designing genetic circuits goes a step further and requires knowledge of features underlying a dynamic response as a result of the binding interactions. Currently, data availability constraints often diminish the impact of ML models in biology, which require large and diverse datasets and become less accurate when applied to situations that significantly differ from those they are trained on [59, 60]. Understanding these limitations is critical for the field to develop and become widely adopted – confidence in models is crucial for their deployment in industrial settings, where inaccurate model predictions can have high time- and financial-cost.

To address these challenges, in this paper we develop a generative model that outputs new genetic circuits when prompted with a target dynamic function. The goal is twofold; we aim to measure how well a generative model can accomplish this synthetic biological design task for the target function of signal adaptation while also gaining insights into the circuit features that are important for determining the level of adaptation as interpreted by the model. Since adaptation has been studied in depth for biological networks, we can compare our expectations against the features the model *actually* utilises. Following this, we further explore whether the generative model can learn evolutionary properties and thus generate circuits that are adaptable and evolvable.

To overcome the limitations of physical experiments and explore a vast number of mutations, we used RNA simulators to create a large dataset of RNA circuits and simulated their dynamics. We then trained a simple conditional variational autoencoder (CVAE) on these circuits, using their adaptation levels as labels. We found that even this small and simple machine learning model could generate new circuits with a desired level of adaptation. Deeper analysis of the model’s embedding space confirmed that the CVAE had successfully captured the specific network motifs that are essential for adaptation [5, 12, 13]. We found that the training dataset can be as small as 2,000 samples to achieve high quality generative results, which has important practical implications for experimental approaches. Finally, by combining prompts for adaptation with those for evolutionary stability, we were able to generate circuits that were not only adaptable but also either evolvable or stable against mutations, with meaningful representations for evolutionary stability.

## 3 Results

### 3.1 Simulated RNA circuits serve as training data for the CVAE

We first construct a digital environment for genetic circuits that enables both dynamical simulation and exploration of the evolutionary landscape. In this framework, a circuit is represented as a network whose nodes are biological components (RNA molecules) and whose edges encode pairwise binding strengths. We focus on RNA molecules because robust physics-based simulators of RNA interactions are widely available. For each RNA pair, interaction strength is quantified as the minimum free binding energy predicted by IntaRNA 2.0 [27] (Figure 1a); intuitively, more negative energies correspond to more stable, and therefore stronger, binding. Individual RNAs are generated by randomly sampling nucleotide sequences from a constrained sequence space (see Methods), and these sequences are then used to compute all pairwise interactions with IntaRNA. To simulate circuit dynamics and assess functional properties such as adaptation, we map these energies to reaction rates using experimental parameters from Na et al. 2013 [31]. Finally, we simulate the dynamics of all circuits, designating one RNA (node 1) as the signal “input” and another (node 3) as the “output”.

**Figure 1.**
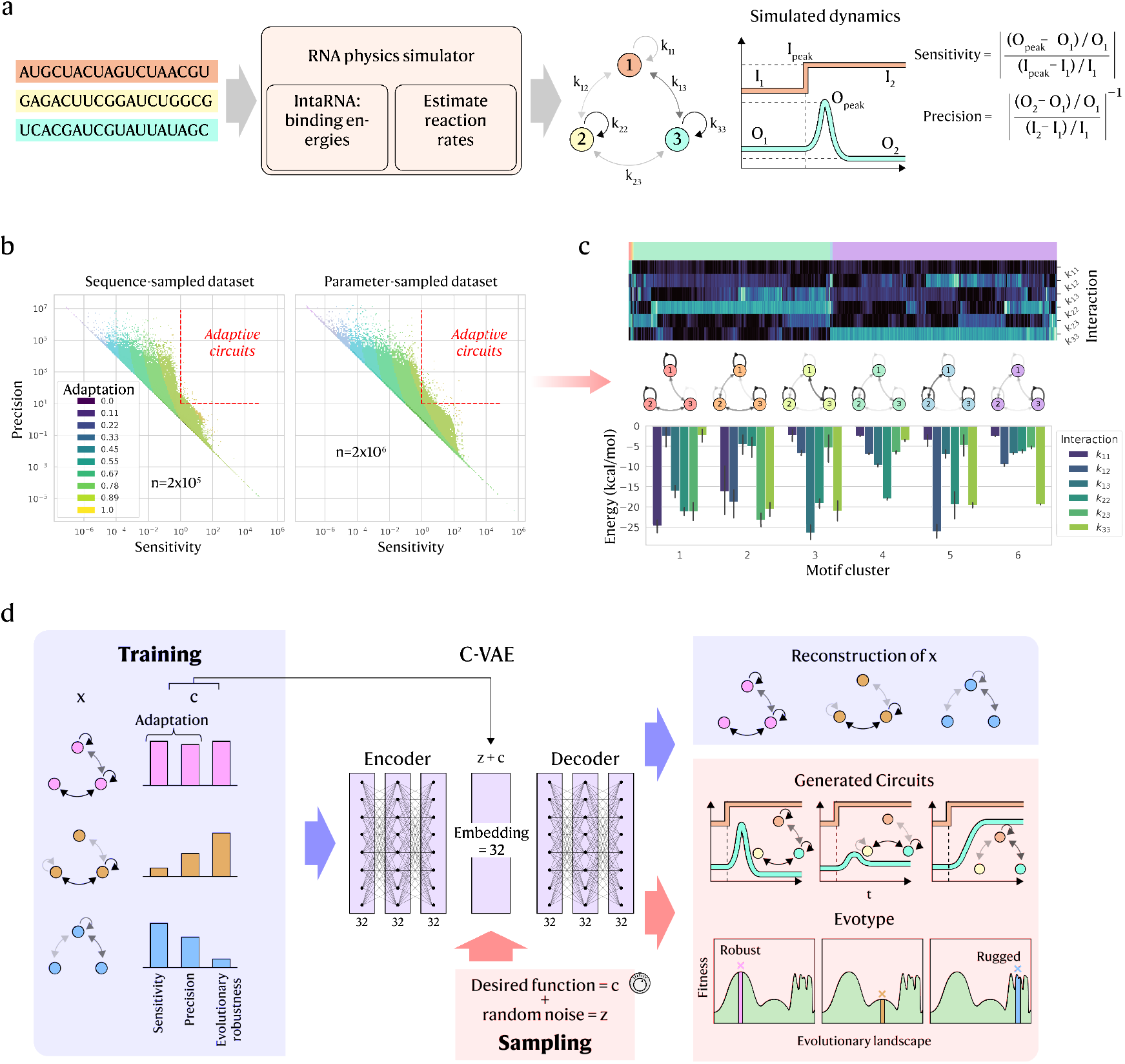
a) Overview of the *in silico* RNA circuit dataset construction: RNA sequences are randomly generated as circuit constituents, then their interactions are simulated with IntaRNA 2.0 and used to estimate reaction rates. From this, the circuit dynamics are simulated and a signal response is recorded to calculate sensitivity and precision, among other functional attributes. b) The custom adaptation function is defined by sensitivity and precision, plotted on a log scale for the training dataset, which is composed of a sequence-sampled (2 × 10^5^ samples) and a parameter-sampled (2×10^6^ samples) dataset. The dashed red line denotes the threshold for adaptation as per [5] (sensitivity *>* 10, precision *>* 1). c) Hierarchical clustering (top) on the interactions of the circuits that fall into the adaptable region interactions identifies several motifs for achieving adaptation. Network diagrams (middle) show the 3-node circuits with the distributions of the interactions present in each motif group (bottom). Interaction arrows in the diagram are the median binding energy of each group. d) Training and sampling the CVAE: For training (blue boxes/arrows), the circuit interactions (vector *x*) are concatenated with the functional properties (vector *c*) and passed through the encoder. The resulting embedding *z* is then concatenated again with *c* and passed through the decoder, which outputs reconstructions of the input circuits. For sampling (red boxes/arrows) a random noise vector instead takes the place of *z* and is concatenated with *c*, which now serves as the prompt, before being passed through the decoder. The generated circuits are simulated to ascertain their dynamic and evolutionary properties.

By simulating each genetic circuit’s dynamics (modeled as RNA concentration trajectories) after adding a step input as the signal, we compute descriptive metrics for every RNA in the circuit, including steady states, overshoot, settling time, signal sensitivity, and signal precision [5]. Sensitivity and precision together quantify adaptation, whereby an adaptable circuit responds strongly to the input (high sensitivity) and subsequently returns to its pre-stimulus level (high precision). We therefore characterise adaptability using the definitions in [5] (illustrated in Figure 1a; formal definitions in Methods) and adopt adaptation as the target function of the circuit (i.e., design objective), as it is well understood yet sufficiently complex to serve as a proof of concept.

In Ma et al. [5], “effective” adaptation is defined by sensitivity ≥10 and precision ≥1 (dotted red boundary in Figure 1b). Such motifs are rare (∼0.01% of random topologies) and enact *integral control*, whereby feedback from an input signal is captured by the circuit components and modulates the output based on the area-under-the-curve between real and target output [61]. Because this binary criterion discards gradations below the thresholds, we instead use a continuous objective that combines sensitivity and precision into a smooth radial field centred at high values and increasing toward the threshold region (shaded in Figure 1b; full definition in Methods). This enables circuit designs to climb smoothly toward the optimum across the design space.

Beyond the adaptation-capable region, the space of RNA sequences that interact *at all* is also sparse. Random sampling mostly yields inert circuits, where components show negligible binding in IntaRNA with minimum free energies near 0. To address this, we supplement the sequence-sampled dataset with a parameter-sampled dataset that draws directly from possible circuit parameters (without simulating pairs of RNA sequences) to assign a minimum free energy to each interaction (Figure 1b, right). The generative model is trained on both datasets, representing each genetic circuit as a numerical matrix of binding strengths.

Since we expect the generative model to recover adaptive motifs, we first distil candidate network motifs from the training data via hierarchical clustering to serve as baselines for comparison (Figure 1c). Prior work on enzyme networks shows that adaptability requires at least one of two motif classes, including buffer/opposer or proportioner/balancer [5, 11–13]. Although RNA circuits interact only via mutual binding (not explicit activation or repression), the same topological rules apply if we view RNA circuits as enzyme networks restricted to mutual repression. In both motif classes, adaptation emerges through integral action, where one or more nodes temporarily accumulate and then release concentration, driving the output back to the initial concentration. This can arise readily in RNA circuits from self-interactions and imbalanced reaction rates (see Supplementary Note 1), so we expect the generative model to exploit such motifs.

For the generative model, we use a variational autoencoder (VAE) as a simple, strong baseline [54, 62], and adopt its conditional form (CVAE) to target specific circuit functions [63, 64]. In training (Figure 1d), the encoder receives the vector *x* concatenated with the label *c*, where *x* encodes the circuit topology via interaction parameters [*k*_11_, *k*_12_, *k*_13_, *k*_22_, *k*_23_, *k*_33_] and *c* is the adaptation metric; both are normalised to [0, 1]. The encoder outputs a 32-dimensional latent representation *z*. Because CVAEs are probabilistic, *z* parameterises a distribution conditioned on *c*, with a KL-divergence term encouraging distinct latent representations for different prompts. The decoder then takes [*z, c*] to reconstruct the original circuit, scored by a reconstruction loss (see Methods). We inject *c* at the latent stage so that the trained decoder can be used alone later for generative purposes. Overall, training organises the latent space so circuits with similar dynamics cluster together, enabling the decoder to sample new circuits that achieve desired functions.

After training, sampling new circuits is straightforward. We supply the generative model with a target function (e.g., a desired adaptation value) and obtain a circuit as a set of binding parameters predicted to meet that target. At sampling (Figure 1d), the encoder is bypassed and the prompt is fed directly to the trained decoder, with random noise used in place of the latent *z* to produce diverse samples. Because the encoder learned a conditional distribution, the decoder remains sensitive to prompts. We then simulate each generated circuit’s dynamics with the same numerical pipeline used in training to evaluate prompt adherence. We next report training results before analysing accuracy and adherence in detail.

### 3.2 The CVAE encodes meaningful representations

For adaptation, the CVAE reconstructs circuits with high accuracy (*R*^2^ = 0.97, Figure 2a) and generates diverse circuits across prompts, despite its simple architecture. We assess model quality based on how well the CVAE reconstructs circuits and how specifically it responds to prompts. While samples need not match a prompt exactly since sampling noise induces variation, the generated distributions should be centred around the prompt.

**Figure 2.**
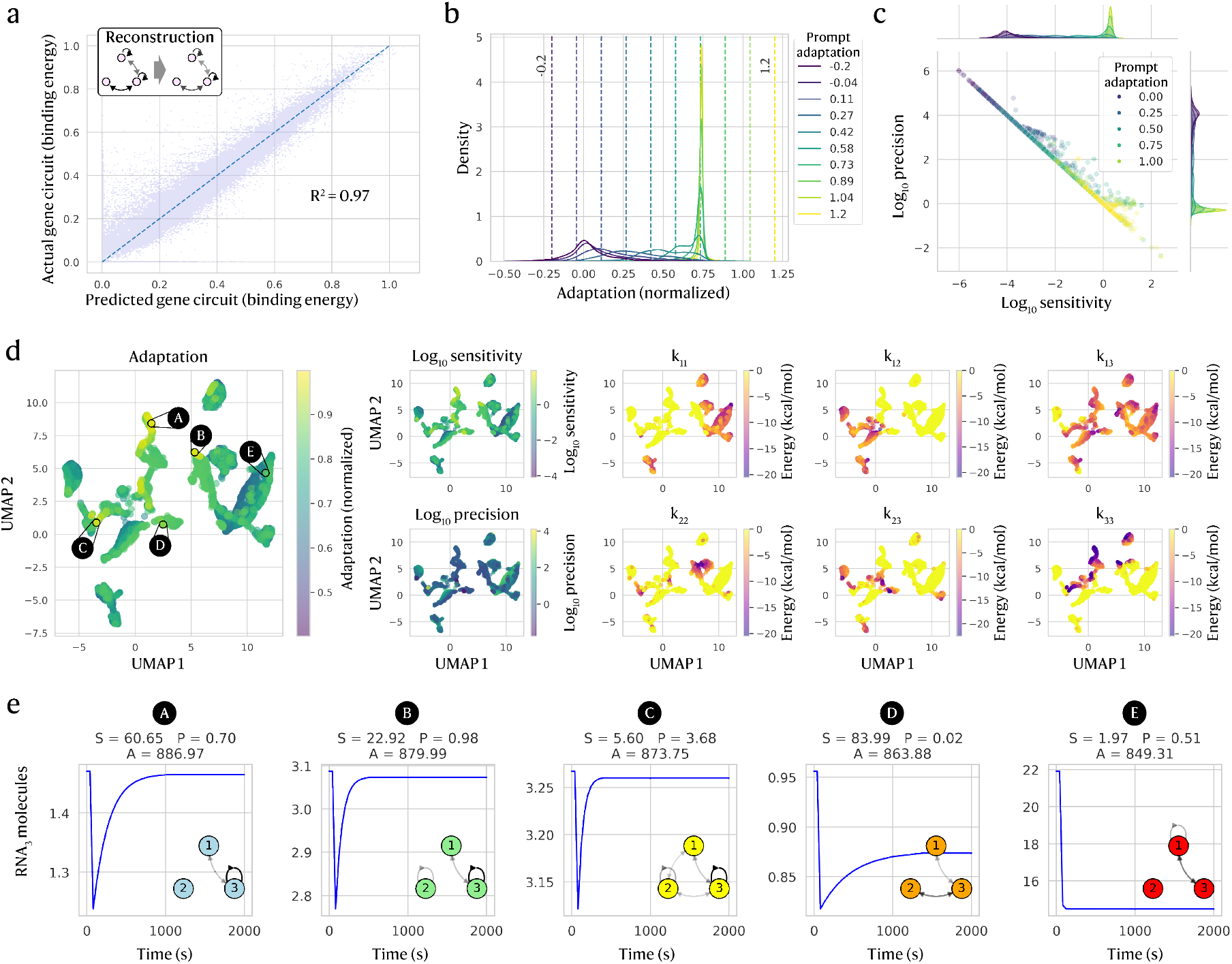
CVAE trained on adaptation generates novel circuits faithfully. a) CVAE-reconstructed circuits vs. true circuit (interactions in terms of minimum free binding energy) shows high accuracy for both training and test data, with an *R*^2^ of 0.97 for test data. b) Kernel density estimation (KDE) distributions of generated circuits (n=10,000) by real adaptation (transformed to the normalised training data range [0, 1]) show high adherence of each distribution to the intended adaptation prompt. Coloured dashed vertical lines correspond to the target prompt. c) Generated circuits show adherence to prompts in both sensitivity and precision, the components of adaptation. d) UMAP of the simulated circuits reveals a latent space structured by adaptation (left). To the right, subplots show how the latent space is structured in terms of sensitivity and precision, as well as how each unique circuit interaction [*k*_11_, *k*_12_, *k*_13_, *k*_22_, *k*_23_, *k*_33_] varies between clusters. e) To illustrate actual dynamic behaviours, 5 circuits with the highest adaptation in each of their clusters (A-E) are picked out from the UMAP plots, with different motifs represented.

To evaluate prompt adherence, we sampled 10,000 circuits across 10 adaptation prompts. Since inputs were normalised to [0, − 1], we can push the generative model to extrapolate outside the outside of the training range with prompts in the range [−0.2, 1.2] (vertical lines in Figure 2b). A weak model would yield similar adaptation distributions for all prompts, while a strong one would peak near each target. Our CVAE produces distinct distributions with peaks near their prompts that increase monotonically with prompt order (Figure 2b) and still generates some circuits beyond the training range, despite sparsity in training data.

Although the prompt-wise distributions peak near their targets, some are broad, especially for lower prompts *<* 0.5, which show long tails (Figure 2b). Plotting sensitivity and precision separately (Figure 2c) shows strong overall alignment with targets but a slight high-end bias, as some circuits prompted with adaptation = 1 reach higher sensitivity/precision than those prompted with *>* 1, suggesting limited extrapolation beyond the training range. We also see a peak at *s* = 1, *p* = 1 for medium–high prompts, likely due to overrepresented weak-response circuits and sparse highly adaptable ones. These patterns reflect biases from both the quantitative objective (our adaptation metric) and training-data imbalance, a common challenge in computational biology [60].

Despite these biases, the CVAE learns meaningful latents that cluster by adaptation level (Figure 2d). We project embeddings with uniform manifold approximation and projection (UMAP) to assess structure. A naive UMAP on raw circuit parameters showed only weak gradients with substantial overlap (Supplementary Figure S1), implying richer representations are needed. In contrast, UMAP of CVAE latents separates circuits by adaptation. Overlaying interaction energies (Figure 2d, right) highlights which parameters drive clustering – *k*_11_ and *k*_33_ are most determining, whereas *k*_13_ and the self-interaction *k*_22_ vary within clusters. Self- and cross-interactions at the input/output nodes (1 and 3) thus primarily set adaptation in the latent space, while interactions with the auxiliary node 2 capture finer differences among adaptation-capable motifs consistent with previous literature [5, 13].

To examine adaptation-capable motifs, we analysed the dynamics and topology of the most adaptable circuit from each latent cluster (black circles in Figure 2d,e). The cluster with circuit A contains highly adaptive circuits, while clusters with B and C show medium–high adaptation and clusters with D and E show medium–low adaptation (see trajectories in Figure 2e). Circuits A–C reach similar adaptation via distinct topologies, whereas D and E respond to the signal but fail to adapt well. UMAPs overlaid with interaction parameters confirm that clusters map to distinct motif classes, indicating that the latent space encodes meaningful diversity.

These cases reveal clear embedding-space trends, showing that the model learns broadly informative representations, though some motif classes remain partially overlapping. Furthermore, the robustness of circuit architectures influences how well the generative model can distinguish it from similar designs. Motifs that are evolutionarily unstable (functional yet on rugged fitness landscapes) sit only a few mutations from non-functionality and are easier to misclassify than robust, constrained motifs. At the same time, it is important that the generative models does not miss these unstable motifs. Next, we probe factors that shape generative performance by varying hyperparameters and prompt formats.

### 3.3 Model parameter optimization and evaluation

Generative model performance acutely depends on many hyperparameters spanning training, optimisation, losses, and architecture, particularly when data are limited. We initially performed broad parameter sweeps to set appropriate ranges (see Methods) before focusing on the most consequential factors, including KL-divergence weight, training dataset size, and the objective function. We compare settings by reconstruction accuracy (*R*^2^) and prompt adherence (Figure 3), evaluating the latter by how distinctly prompt-wise distributions separate across different prompts.

**Figure 3.**
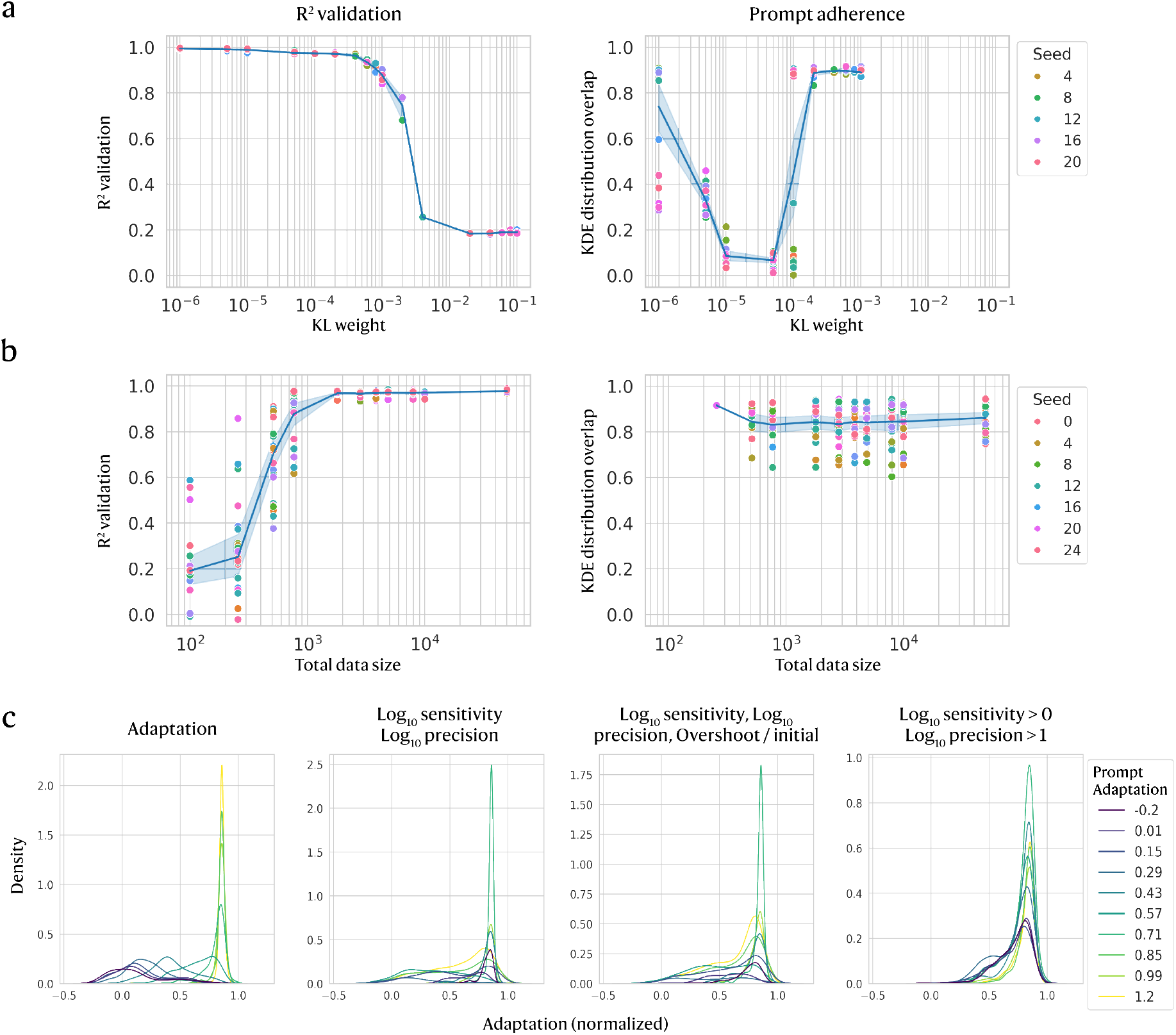
Generative model training with varying KL weight, dataset size, and objective function. a) The weight controlling KL divergence loss is varied and compared by model accuracy (*R*^2^) and prompt adherence, which is assessed for the simulated dynamic function adaptation by measuring the average overlap between the KDE distribution of a prompt and all other prompt distributions. For each value, 20 different seeds are used for model initialisation. b) Similarly to a), the accuracy and prompt adherence are assessed across training dataset sizes, with 24 seeds used for model initialisation. c) To examine how using alternative representations of adaptation affects model training, we compare the KDE prompt distributions for the best-performing model (out of 60 initialised with different seeds) for each of the 3 new objective functions, [log_10_ *S*, log_10_ *P*], [log_10_ *S >* 0, log_10_ *P >* 1], and [log_10_ *S*, log_10_ *P*, overshoot/initial].

A core feature of VAEs, the KL divergence weight sets how strongly we penalise similar embeddings across different prompts. Parameter sweeps (Figure 3a) reveal a narrow workable range; model accuracy collapses near 10^*−*3^, while prompt adherence is high for weights 10^*−*5^–10^*−*4^ (low overlap among prompt-wise distributions). We then tested data requirements (Figure 3b) and found that although the sequence-sampled and parameter-grid datasets respectively contain ∼100,000 and ∼1,000,000 samples, as few as ~ 1,500 training points achieved high *R*^2^ without degrading prompt adherence. In particular, prompt adherence varies with the random initialisation seed, which presents considerable implications for using generative models in experimental settings.

Beyond training parameters, we find that the choice of objective function strongly shapes learning. Our custom adaptation metric collapses sensitivity and precision into a single value, masking different (*S, P*) combinations. We therefore compared this to multi-parameter objectives that treat metrics separately (Figure 3c): a continuous target [log_10_ *S*, log_10_ *P*]; a binary categorisation [log_10_ *S >* 0, log_10_ *P >* 1] [5]; and a three-metric target [log_10_ *S*, log_10_ *P*, overshoot*/*initial], motivated by the link between overshoot and adaptability. For each objective, we trained 60 models with different seeds and generated 3,000 circuits using prompts in [−0.2, 1.2]. For the binary case, we also tested intermediate prompts to assess whether the generative model implicitly learned beyond just categorical labels. To limit combinatorial explosion among prompts, we used 10 prompts for the unified adaptation objective and 5 prompts per dimension for multi-objectives (e.g., 5^2^ = 25 for [log_10_ *S*, log_10_ *P*] and 5^3^ = 125 for the three-metric target).

Because prompting differs across objectives, we cannot compare models with the overlap of prompted distributions. Instead, we map each multi-objective prompt to an equivalent adaptation value (combining sensitivity and precision) and compare that to the true adaptation of generated circuits (Figure 3c). For [log_10_ *S*, log_10_ *P*], distributions are separated but misordered relative to their targets, likely because the precision part of the prompt influences the output disproportionately (Supplementary Figure S2). Adding overshoot ([log_10_ *S*, log_10_ *P*, overshoot*/*initial]) does not substantially improve prompt adherence (3c), although the sensitivity part of the prompt shows marginally better alignment to actual sensitivity (Supplementary Figure S2). For the binary objective [log_10_ *S >* 0, log_10_ *P >* 1], the gradation of the prompt-wise distributions indicates that the generative model learned to interpolate beyond just categorical values. Although distributions peak further from targets and the overall adaptation range is narrower than the continuous objectives, prompt-wise distributions are ordered and distinct, showing that even a simple binary objective can outperform more complex ones.

Compared with a single adaptation objective, multi-objective training struggled to learn multiple high-dimensional conditional distributions and to follow prompts. Redefining the objective function with a custom scalar function (adaptation) or simplified formulation (e.g., binary) often produced better generated outputs than prompting with raw metrics. Simple objectives are therefore a good starting point before introducing more complex custom functions. With balanced data, even small training datasets can yield strong models, which is feasible for circuits with few interacting components as investigated here. Next, we examine a different case of multi-objective training with the wholly distinct objective functions of adaptation and evolutionary ruggedness.

### 3.4 Extending the CVAE to include evolutionary properties

Having shown that the CVAE can generate adaptation-targeted circuits even with limited data or binary labels, we next incorporate evolutionary stability into design. We estimate ruggedness by perturbing a circuit and measuring the resulting change in function. This can be done by (1) mutating RNA sequences and recomputing binding energies or (2) directly perturbing the interaction energies between RNAs. We opt for (2), since sequence-level details are not provided to the CVAE. Our ruggedness metric slightly perturbs each unique interaction, re-simulates the dynamics, and aggregates the resulting changes in adaptation across all perturbations.

Since multiple genetic circuit topologies can yield the same adaptation, a fixed adaptation level can lie on rugged or smooth regions of the evolutionary landscape. Nevertheless, some correlations arise from the definition of ruggedness ruggedness, which perturbs each unique interaction and combines the change in adaptation between original and perturbed circuits (Equations 6, 7). Because adaptation weights sensitivity slightly more than precision, ruggedness is biased with perturbations that mainly affect sensitivity. Consequently, ruggedness decreases mildly as adaptation increases (Figure 4a), while sensitivity and precision individually are largely flat versus ruggedness (Supplementary Figure S3). A further bias comes from the logarithmic transform used to temper very high adaptation values, which also expands the low-ruggedness region so that roughly half of the [0, 1] normalised range maps to negligible true ruggedness *<* 0.01. As a result, most circuits are compressed into a narrower effective range starting at a normalised ruggedness of 0.5, especially affecting adaptable circuits that only make up a small proportion of the training data (Figure 4a).

**Figure 4.**
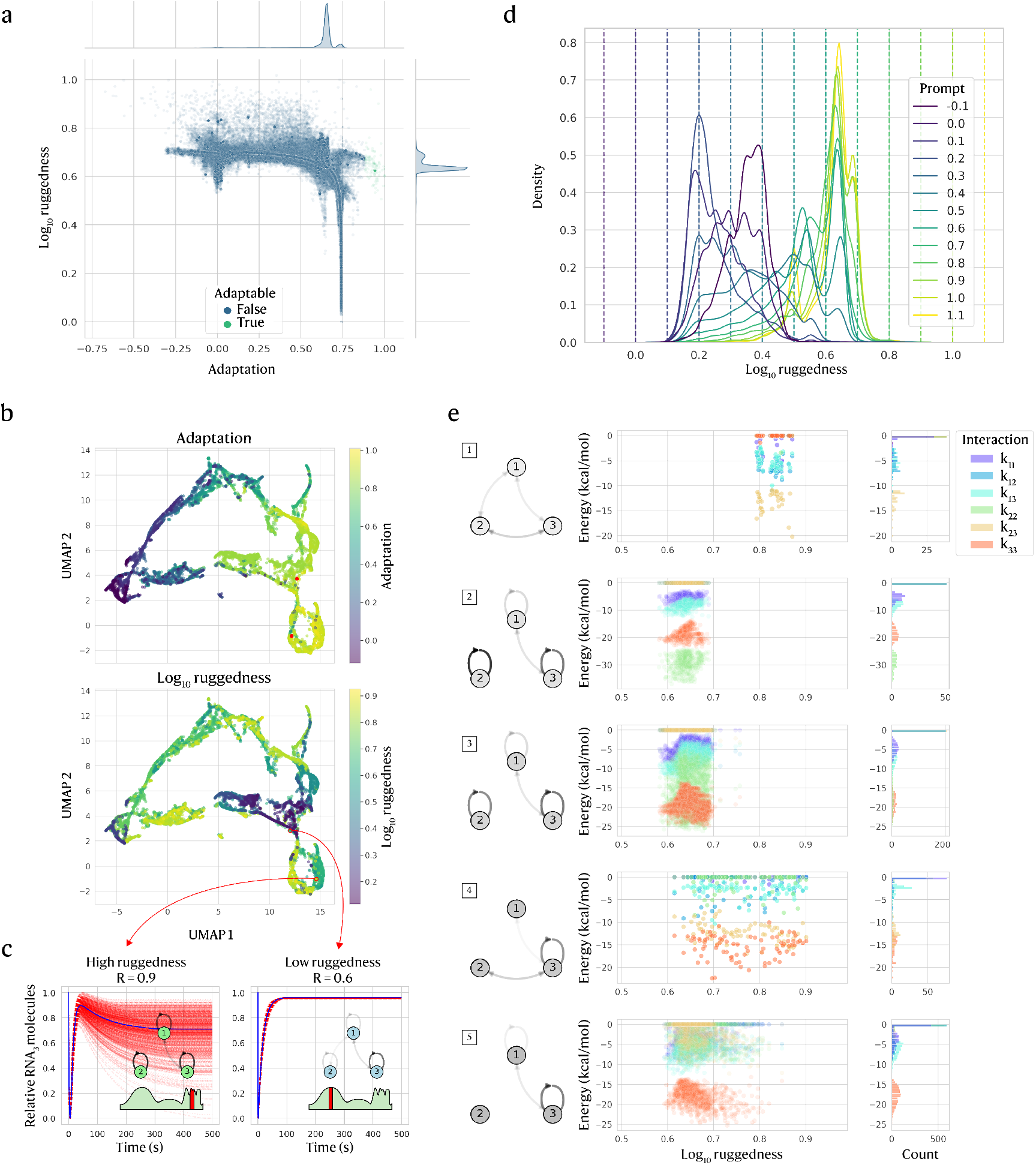
Adaptation and evolutionary ruggedness are effective prompts for generating circuits. a) Adaptation correlates slightly with log_10_ ruggedness, shown for training data before pre-processing but normalised to [0, 1] (samples outside of this range are later excluded). b) UMAP of generated circuits demonstrates that the latent space is structured by both normalised adaptation (top) and ruggedness (bottom). c) Two circuits chosen from a high-adaptation region with high (left) and low (right) ruggedness are plotted by output RNA (blue) against 1,000 mutated versions (dashed red), normalised to a [0, 1] range to show relative changes. Network diagrams show the original circuit topology and an illustration of the evolutionary landscape. d) KDE prompt-wise distributions of ruggedness prompts (prompted adaptation = 1) are distinct and adhere well to most ruggedness prompts. e) Hierarchical clustering of interactions in adaptable circuits (*S >* 1, *P >* 10) identifies 5 motifs that also differ in ruggedness, with network diagrams (left) representing median circuit interactions.

Beyond simulating the ruggedness metric, inferring stability from topology alone is difficult. Nevertheless, the CVAE’s latent space (Figure 4b) clusters by both adaptation and evolutionary ruggedness, with relatively high-adaptation regions splitting into high-vs. low/medium-ruggedness subregions. We illustrate this with two CVAE-generated circuits that are both largely adaptive but differ in ruggedness (Figure 4c). We plot the output RNA (blue) alongside perturbed variants (dashed red), normalised to [0, 1] to emphasise relative changes. The high-ruggedness circuit (left) shows large trajectory shifts after perturbation, whereas variants of the low-ruggedness circuit (right) closely track the original, preserving sensitivity and precision (see Supplementary Note 2 for a mechanistic account).

The structured latent space suggests that joint prompting for adaptation and ruggedness is coherent, so we test adherence to combined prompts. Across many pairs of prompts, generated circuits concentrate near their targets. At adaptation prompt = 1, ruggedness distributions are distinct for ruggedness prompts (Figure 4d), although extremes prove more challenging – very low ruggedness prompts (*<* 0.2) rebound to higher true ruggedness, while low–medium ruggedness prompts peak near the intended level. High prompts (*>* 0.7) yield narrow distributions that saturate around ∼0.7, with a small tail to ∼0.9. Because ruggedness combines adaptation changes over all perturbed interactions, high ruggedness arises more readily when many topological links are vulnerable. These effects are less pronounced at other adaptation prompts with more training data (Supplementary Figure S4). Despite data limits, the model still produces low-ruggedness circuits at high adaptation, indicating it learns useful combinations of objectives from the data.

As exemplified by the adaptable pair in Figure 4c, adaptable motifs can be either mutation-prone or mutation-resistant. The CVAE’s latent space organises circuits by ruggedness, suggesting it captures topology links to evolutionary properties. Using hierarchical clustering, we extracted motifs from CVAE-generated adaptive circuits and related them to ruggedness (Figure 4e; Supplementary Figure S5). Cluster 1 is high-ruggedness (evolvable), clusters 2–3 are low-ruggedness (stable), and clusters 4–5 span intermediate levels (flexible). As stability increases, allowable interaction-energy ranges narrow (e.g., motifs 2, 3, and 5), consistent with core motif “constellations” that become more evolvable when pushed toward their boundaries in high-dimensional design space (see Supplementary Note 3). The CVAE reflects this, as ruggedness prompts are higher in genetic circuits that deviate from their most stable settings (radar charts in Supplementary Figure S6). Thus, the CVAE can generate both evolvable and stable circuits while maintaining adaptation.

## 4 Discussion

The challenges facing engineering biology are complex, multi-faceted and well-suited for machine learning approaches. Genotype-phenotype datasets that combine underlying sequences with behavioural features such as dynamic functions or metabolic properties are increasingly being pursued experimentally to fill the data gap for training high-quality models [65–71]. However, more data is not always the answer – understanding the limitations of ML models, especially generative approaches, is an important step in order to make them trustworthy, deployable, and competitive with other discovery methods [60, 72, 73].

In this paper, we present a novel generative approach for realising genetic circuits that optimise both functional and evolutionary properties, and show that even simple generative models can identify key features that recapitulate known adaptation motifs (as identified in [5, 12, 13]). Even without providing full time-series data (as in [54]), summary metrics like sensitivity and precision (even as binary labels) are enough to train our model and output diverse designs. Many generative approaches designed sequences of individual biological parts [48, 50, 74–79] while other ML approaches predicted optimal circuit composition and culture conditions [40, 68, 80, 81], so it is plausible that multiple sequences could be generated for a desired dynamic behaviour. Although our simulated RNA circuits dataset makes many assumptions and approximations, this work serves as an important proof of concept for utilising generative models in genetic circuit design.

Our work also highlights many of the challenges and tradeoffs faced when quantitatively defining an objective for circuit behaviour. Our custom scalar for adaptation improved accuracy and prompt adherence but obscures information about mid-range combinations of sensitivity and precision. However, treating these components as separate targets reduced prompt adherence; adding more objectives (e.g. overshoot) skewed the training distribution and hindered conditional learning. One possible improvement could be encoding raw metrics in alternative ways (e.g. one-hot). The joint objective (adaptation and ruggedness) likewise suffered from poor training representation of some combinations, despite our large *in silico* dataset. While specificity was reduced at extreme prompts, the model performed well even for some data-poor combinations, suggesting latents learned in one range can generalise more broadly.

We attempted to improve embedding quality and sought a latent space structured even more by circuit function using several strategies. After tuning the KL weight, architectural tweaks (e.g., embedding size) yielded little improvement, so we added a contrastive loss to pull similar samples together and push dissimilar ones apart [82, 83]. This neither improved prompt adherence nor showed better organisation of the latent space (Supplementary Figure S7). Other contrastive implementations [84–86] or alternative latent encodings [87] may help, as well as VAE variants aimed at disentanglement [88–90]. Some issues are intrinsic to VAEs, including poor approximations of heavy-tailed distributions [91] and sensitivity to weight initialisation seeds [92–95], which can determine whether a model responds to prompts at all. We found no reliable heuristic to predict prompt adherence without simulating generated circuits, which is analogous to costly lab tests and illustrates practical challenges in deploying VAE models. Alternative models, such as hybrid graph networks that generalise well and can handle network inputs, may address these limitations [96].

Nevertheless, the CVAE generally performed well, especially considering the sparsity of adaptable circuits and retention of accuracy training on just over 1000 samples, which could be achievable in the lab. For practical applications with expensive data collection and optimisation processes often relying on scientific intuition, flexible ML models can be invaluable in identifying good initial designs for testing, and support existing model-driven algorithms like Bayesian optimisation (BO) and other iterative/active learning approaches that iterate on these designs. These are already heavily used in industry [97–100] and show promise for synthetic biology [25, 77, 101, 102]. Furthermore, many ML models can estimate their own uncertainty about a prediction and utilise diverse inputs such as sequences or images, presenting a compelling upgrade to existing methods that are limited to categorical or continuous inputs. Advanced models like transformers pose an especially suitable option for integrating sequence-level information (such as mutation type and location) with more abstract circuit features (such as switches or logic gates) [103] and thus predicting evolutionary properties from sequences directly [49, 104].

Beyond existing models, we argue that there is an opportunity to further develop machine learning tools and architectures such as hypergraph models, biologically inspired neural networks, self-supervised learning, or hybrids for synthetic biological tasks [96, 105–109]. Novel architectures better geared to operate at multiple levels of abstraction than VAEs could reduce more complex biological networks into overarching motifs, such as large adaptable biochemical pathways [13] and genetic circuits themselves [103]. Crucially, models that can predict biological interactions beyond RNA-RNA interactions, like protein-protein or protein-DNA, have transformative potential for synthetic biology by integrating sequence-level approaches with functional [79, 110–114] and evolution-aware [104] design. This could broaden our approach from simulated RNA circuits to interacting genes and proteins and allow exploration of new dynamics-predicting models. Ultimately, many exciting innovations on the horizon promise to further tackle the sequence-to-function challenge and enable evolution to become a more predictable aspect of engineering biology.

## 5 Methods

### 5.1 Digital genetic circuit environment

For the sequence-sampled dataset, we first generate a set of *n* = 3 RNA sequences of length *m* = 20 by sampling from a distribution of nucleotides. Their probabilities are equivalent to the occurrence of each nucleotide in *Escherichia coli* cells, with the following proportions: [A: 0.2451, C: 0.2458, G: 0.2622, U: 0.2469] [115]. We then take this collection of RNAs and simulate the binding energies for all paired combinations of sequences as query and target with the IntaRNA interaction simulator using default settings [27]. For each RNA pair including self-interactions, we use the most probable predicted minimum free energy (*kcal/mol*) and convert this to a rate of binding. We use the Gibbs free energy equation to convert from binding energy to ratio of association and dissociation rates:

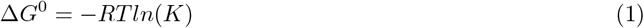

where Δ*G*^0^ is the minimum free binding energy at equilibrium, *R* is the gas constant, *T* is the temperature in Kelvin, and *K* is the equilibrium constant. This equilibrium constant is a ratio of the forward and reverse binding rates *k*_*f*_ and *k*_*r*_, respectively (also known as the association rate *k*_*a*_ and dissociation rate *k*_*d*_). Assuming that the forward rate for RNA binding is very fast [116–118], we fix *k*_*f*_ = 1*x*10^6^ and find the reverse rate *k*_*r*_ from the simulated binding energy and the equilibrium constant *K*. In practice, the *K* calculated directly from the Gibbs free energy equation 1 does not align well with observed RNA binding, for example between small RNA (sRNA) and messenger RNA (mRNA) in RNA circuits controlling green fluorescent protein (GFP) fluorescence [31]. We therefore redefine the conversion from binding free energy to the equilibrium constant using an approximate parameterisation of the observations from Na et al. [31] to get the following expression:

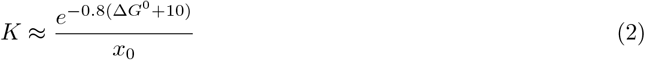

where *x*_0_ is the initial concentration of RNA reactants. The expression stems from the parameterisation of the relative GFP fluorescence measured by Na et al. across different sRNA’s of varying binding energy strength and indicates the amount of binding between the GFP mRNA and an sRNA library acting as inhibitors of mRNA transcription. From this reparameterisation of *K* and with the forward rate constant *k*_*f*_, we can approximate the rates of reaction for simulated genetic RNA circuits. From these rates, we construct a system of ordinary differential equations (ODEs) and simulate the dynamics with the solver library diffrax [119] using the numerical solvers Tsit5 (Tsitouras’ 5/4 method, 5th order explicit Runge-Kutta) and Dopri5 (Dormand-Prince’s 5/4 method) with adaptive step sizing.

This way, the parameter-sampled dataset is constructed by sampling the minimum free binding energy Δ*G* directly. For each unique interaction, we sample a value from a uniform distribution with the range [0, 1] and scale the interactions to the range [-30, 0], to represent a realistic range of kcal/mol that an RNA sequence with 20 base pairs could take. These energies are then transformed into equilibrium constants and to reaction rates in the same way as for the sequence-sampled dataset. Because of this fast energy construction step, the computationally intensive RNA sequence interaction prediction simulation can be avoided and a greater diversity and volume of circuits can be sampled.

We simulated the signal response dynamics in two phases. First we simulated the steady states of all free and bound RNAs for all circuits and used these as the initial state for calculating sensitivity and precision. We then perturbed them with a spike (step input) in the input RNA (node 1 / *RNA*_1_). We simulated the dynamics following the settling of this spike into new steady states and used these as the final states for adaptation calculations. The default parameters relating to the signal response were kept fixed across all simulations. These molecular parameters are summarised below:

The code used to generate the dataset can be found in the GitHub repository https://github.com/olive004/genetic_circuit_generator.

**Table 1:** Molecular Parameters for Dynamic Simulation.

| Parameter | Value |
| --- | --- |
| Average mRNA per cell ($mRNA$) | 100 |
| Cell doubling time ($s$) | 1200 |
| Starting copy numbers ($mRNA$) | 100 |
| Number of mRNA per circuit | 3 |
| Sequence length per mRNA | 20 |
| Signal (input) mRNA | $RNA_1$ |
| Output mRNA | $RNA_3$ |
| Amount of impulse signal added to steady state | $2x$ steady state $RNA_1$ |
| Degradation rate ($mRNA\ s^{-1}$) | 0.01175 |
| Association binding rate ($mol^{-1}\ s^{-1}$) | $1x10^6$ |
| Association binding rate ($mRNA^{-1}\ s^{-1}$) | $1.50958097 \times 10^{-3}$ |

### 5.2 Defining sensitivity and precision

More formally, sensitivity measures how responsive the output is to the input signal, defined as the maximum change in the output relative to the change in the input (Equation 3). Conversely, precision measures how closely the output returns to its initial value before the input signal was added, defined as the inverse of the relative change in the output compared to the change in the input (Equation 4):

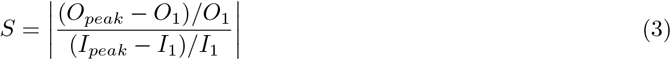

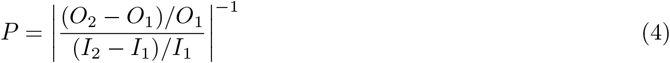

where *S* is sensitivity, *P* is precision, *O* and *I* are the output and input species concentrations (here in molecules / cell), the subscript 1 denotes the initial concentration, the subscript 2 denotes the final concentration, and the subscript *peak* denotes the peak concentration (e.g. the maximum or minimum achieved during a simulation period, depending on the direction of change), see figure 1a (right) for an illustration of these features.

### 5.3 Defining adaptation to signal inputs

Sensitivity and precision are metrics used to assess whether a system is able to adapt to a signal [5] and have been defined in Equations 3 and 4. However, there is no continuous metric for adaptation for biological networks beyond the binary definition that circuits with a sensitivity *>* 1 and a precision *>* 10. For training a model and optimising circuits towards a single objective, we constructed a metric for adaptation that is positive within the region where circuits function, encompassing the ranges [10^*−*7^, 10^2^] for sensitivity and [10^*−*2^, 10^7^] for precision. The adaptation metric is defined as a concentric gradient increasing towards the binary threshold as the following:

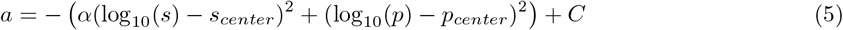

where *s* and *p* are sensitivity and precision, *s*_*center*_ = 7 and *p*_*center*_ = 7.5 are their respective concentric centers, *α* = 3 is the adjustable extra weight given to sensitivity, and *C* = 1000 is a constant to ensure the metric is positive in the operational region. Making the metric positive facilitates calculating the loss function across all sensitivity and precision values so that the loss always has the same sign. The circuits also always move in the correct direction across both dimensions thanks to the the centers of sensitivity and precision, which allow adaptation to peak at sensitivity = 10^7^ and precision = 10^7^.5 well within the adaptation threshold region. The precision peak is slightly higher to reflect that the binary boundary also has a higher precision. This alone was not enough to bias optimisation towards more sensitive circuits and high precision values easily dominated, so we added the adjustable weight *α* to boost sensitivity slightly more. Values that go to infinity due to sensitivity or precision having divisions by zero are set to the Python JAX *NaN* value.

### 5.4 Defining evolutionary ruggedness

To compare circuits in terms of their evolutionary properties, we perturb their parameters by a small fixed value and resimulate the dynamics of the mutated circuit. Circuits can be defined by the number of unique interactions between nodes, which for a 3-node circuit results in 6 unique interactions [*k*_11_, *k*_12_, *k*_13_, *k*_22_, *k*_23_, *k*_33_]. The ruggedness of the evolutionary landscape on which a circuit sits can be seen as the magnitude of its gradient with respect to each interaction. We therefore calculate ruggedness by perturbing each interaction individually and then summing up the squared changes in adaptation relative to the perturbation applied across all mutated versions of a circuit. This can look like the following equation for each circuit:

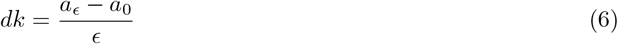

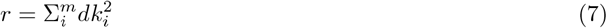

where *dk* is the change in adaptation per perturbation *ϵ*, which is set to *≈* 1 kcal/mol binding energy, *a*_*ϵ*_ and *a*_0_ are the perturbed and initial adaptation values respectively, and the ruggedness *r* is a sum of squares across all *m* = 6 unique interactions *dk*_*i*_. For the perturbed circuits in Figure 4c, the red traces are mutated versions of the original that have had all of their interactions perturbed by a small, random amount (normally distributed around 0, with a standard deviation of 10% of the strongest binding energy in the training data).

### 5.5 CVAE model architecture

We implemented the conditional VAE model with the Python packages JAX [120] and Haiku [121]. The general VAE architecture was kept the same as the default model [63, 121] composed of an encoder and a decoder, with the main changes occurring in the number and size of layers. Both the encoder and decoder are a stack of 3 linear layers of size 32 with leaky ReLU (rectified linear unit) activations and He normal (Kaiming normal) weight initialisation. While encoders and decoders typically do not have same-sized layers, an initial hyperparameter sweep changing the layer sizes did not yield significantly better results and the same-sized layers were sufficient for our purposes. We do acknowledge that this would be a source of improvement for optimising model architecture, especially for models deployed beyond simulated genetic circuits. The CVAE model additionally has two layers for transforming the encoder output *h* into the probabilistic parameters *µ* and *logvar*, which are combined to determine the embedding space in a reparameterisation step:

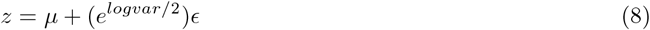

where *z* is the embedding, *µ* is the mean or center of the distribution, *logvar* is the log variance of the distribution, and *ϵ* is random, normally-distributed noise that can be added to provide sample diversity or stochasticity during training for better model robustness.

### 5.6 CVAE model training

#### 5.6.1 Training data

The model was trained with the circuits and their dynamic function of adaptation as input. The circuit input *x* has dimensions [*n, m*] and the input conditional variable *c* has dimensions [*n, c*_*n*_], where *n* is the total number of training samples (eg. *≈* 1*x*10^6^, *m* is the number of unique interactions per circuit (6), and *c*_*n*_ is the number of objectives part of the conditional variable. For adaptation, this would be 1, while using both sensitivity and precision as the conditional variables is also possible and makes *c*_*n*_ = 2. Within the CVAE, the latent variable *z* representing the embedding space has dimensions [*n, H* + *c*_*n*_], with *H* representing the size of the hidden variables *h* and *z*. The conditional variable *c* is concatenated with *z* before being passed to the decoder, hence the dimensions of *z* also depend on *c*_*n*_. The output *y* is just a reconstruction of the circuits, so has the same dimensions as *x*. At the model inference stage where circuits are generated, a noise vector of dimensions [*n, H*] is concatenated as a stand-in for *z* with the prompt *c*.

#### 5.6.2 Data preparation

The inputs *x* and *c* are prepared by filtering and normalising the training data. Circuits with sensitivity and precision with NaN values are removed, along with duplicate circuits and those with exceedingly slow response times that make calculating adaptation difficult. We further normalise circuits with a robust scaling transformation and with a min-max scale to make sure they are in a [0, 1] range. The robust scaling transforms features by subtracting the median from the training data and then dividing by the interquartile range, eg. the range between the 25th and 75th percentile in the data. For the binding energies represented by *x*, we add a negative multiplier to align high model outputs with strong binding interactions. These filtering and normalisation steps stabilise the training process.

#### 5.6.3 Training process

We train the model by optimising its weights with gradient descent over a set number of epochs. At the beginning of every epoch, we shuffle the order of the training data and split it into batches. In each training step, the batches are run through the model iteratively to get a predicted output that is used to calculate the loss and gradients. Since the model outputs reconstructions of the input circuits, a mean square error (MSE) loss is applied initially to penalise differences between the input *x* and the predicted output *y*. For validation accuracy, we calculate how many samples the model reconstructed are within an error bound of 0.1 (out of the [0, 1] range). While we experimented with an L2 loss to ensure that weights remained within a reasonable region, this did not have a significant effect on model accuracy or training stability, so we stopped using it. We next apply a Kullback-Leibler (KL) divergence loss, calculated as the mean of the Gaussian KL divergence loss of the following form [62]:

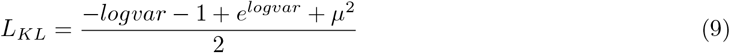

where the *µ* and *logvar* are output by the VAE as the parameterisation of its embeddings. This KL divergence loss is averaged together across all samples and multiplied by a weighting factor, which we explore in Figure 3.

#### 5.6.4 Contrastive loss

Following this, we also tried applying a contrastive loss to improve the quality of latent space clustering, but this did not significantly improve outcomes, as discussed previously (Supplementary Figure S7). We nevertheless report our methods for applying contrastive loss in this context, as there is still reason to believe this could be useful in the future. Contrastive loss typically punishes embeddings that are similar but for which the labels are actually different. Normally, a pair of “same” and “different” samples are compared to an anchor sample based on their membership of a particular category [82, 83]. A batch of similar and dissimilar samples may even be supplied. However, since in our data we are mostly dealing with continuous labels in the form of the adaptation metric, we calculate the distance between all pairs of normalised conditional labels in a batch using a dot product. We then subtract a similarity threshold value to re-center the distance as either positive (similar) or negative (dissimilar). This is then multiplied by the negative of the similarity of the encoder embeddings *h*, which is also calculated with a dot product and scaled by the temperature factor. For a threshold of 0.9, a normalised distance of 0.95 would decrease the loss, while a distance of 0.85 would contribute positively to the loss, depending on the similarity between the embeddings. This can be summarised as the following:

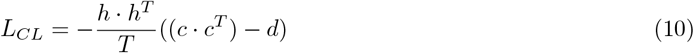

where *T* is the temperature and *d* is the threshold similarity distance. The diagonal of this is also subtracted in order to eliminate counting self-similarities. Improvements to this approach could include simplifying the comparison between samples, for example by comparing pairs of samples to an anchor instead of all samples in the batch, or adjusting the threshold adaptation value at which circuits are considered similar.

#### 5.6.5 Optimisation and hyperparameters

After summing together all relevant losses, we use an Adam optimiser to calculate the weight updates from the gradients [122]. After 20 warm-up epochs in which the optimiser slowly ramps up the learning rate from 0 to 1*x*10^*−*^3 and then sets out on a cosine decay schedule until 2000 epochs are reached. The training loop can stop early if a validation accuracy of over 0.98 is reached or if there is no improvement in the model outputs for 500 epochs. This helps to prevent a model from overfitting on the dataset.

The training process is governed by many hyperparameters, which we determined through automated hyper-parameter tuning followed by sweeps of specific parameters. We used the neural network intelligence (NNI) package [**nni2021**] to find a good starting point and ensure that good parameter combinations are not being missed out on. We quickly found working values for specific variables that had a more noticeable initial effect on model training, such as the learning rate, the validation accuracy threshold, and whether to bin the conditional variable into categories. While categorical labels performed similarly well in terms of accuracy and prompt adherence as continuous adaptation labels, we saw no reason not to favour the continuous labels. More experimentation in this area may be warranted depending on the particular target circuit function.

### 5.7 Model evaluation

While a CVAE can attain high accuracy on reconstructing inputs, this does not automatically mean adherence to prompts in the inference phase. Initially, we used the set of metrics commonly employed in generative AI literature, namely precision, recall, and the combined F1 score [123]. Generally, precision measures the accuracy of positive predictions, while recall measures a model’s ability to find all actual positive instances. In this context, precision is the proportion of circuits generated for a particular prompt that actually fulfill that prompt (within a threshold bound). Recall measures what proportion of circuits that match one of the prompts were actually generated according to that prompt. For example, high adaptation circuits that were generated with a different prompt would be considered missed by the model. The F1 score combined these together to highlight the tradeoff between them in the following way:

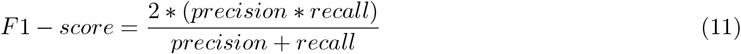

The main reason using these metrics for prompt adherence was difficult was due to the continuous nature of our prompts, meaning that the threshold bounds within which a circuit is considered to adhere to its prompt played a significant role in assessing precision, recall, and F1 score.

While these metrics provided some information about prompt adherence, we constructed a simpler, more heuristic area-under-the-curve metric based on the overlap between distributions of circuits generated with different prompts. We fitted a Gaussian kernel density estimate (KDE) to each prompted collection of generated circuits to create a smooth distribution of their functional properties (e.g. adaptation). We then compare prompt distributions by evaluating the KDEs at 1000 points and summing the number of overlapping points. Overlaps from KDEs are on a [0, 1] scale, with 0 being no overlap and 1 being complete overlap. For each prompt distribution, we use the average of its overlap with all other prompt distributions and finally take the average of these averages. This metric is mostly a heuristic, since we do not account for the number of prompt distributions used to calculate the average area-under-the-curve.

## 6 Data Availability

Data is available on Zenodo at the DOI 10.5281/zenodo.17194085.

## 7 Code Availability

All code can be found in the GitHub repositories https://github.com/olive004/genetic_circuit_generator and https://github.com/olive004/EvoScaper.

## 8 Acknowledgements

This work is funded by the Department of Engineering Science, University of Oxford, and Wadham College, University of Oxford.

## 9 Author information

### 9.2 Contributions

O.G. implemented the algorithms and wrote the original draft. Both authors conceived the idea, developed theory and methods, and contributed to data analysis, discussions, and manuscript preparation.

## 10 Supplementary information

