## Supplementary Information for "Generative design of synthetic gene circuits for functional and evolutionary properties"

### Supplementary information: Sequence-aware computational approaches for generative design of synthetic gene circuits

Olivia Gallup

August 2025

#### 1 Note 1: Adaptation motifs

More intuitively, the buffer and proportioner motifs differ mainly in terms of how directly the signal is relayed to the output and how the output regains its original state. The buffer motif can be enacted in several ways, for example when the output node 3 only interacts with auxiliary node 2 and with itself ($k_{23}$ and $k_{33}$) and relies on node 2 to buffer the signal from input node 1 (i.e., soak up and slowly release the signal so that the output reacts prior to steadying out again). The buffer mechanism can also be enacted by output node 3 if the correct balance between its interactions with node 1 and with itself is struck ($k_{13}$ and $k_{33}$), which is more stable if node 2 is self-binding to interfere less (since some background binding exists even when interaction energy is effectively zero). The proportioner motif gets activated in proportion to the signal and exerts an opposite force on the output, which can be enacted by node 2 if it is not interacting with node 3 directly and still soaks up some of the signal. We thus also expect that the model will identify interactions with node 2 as the determining factor for motifs within adaptation-capable circuits.

#### 2 Note 2: Adaptable circuits with different ruggedness

The topologies of the two circuits are quite similar – both have a strong output node 3 self-interaction ($k_{33}$) and an interaction between RNA nodes 2 and 3 ($k_{23}$). However, the more rugged circuit has a strong interaction between the input and output nodes ($k_{13}$) and relies more on the interaction between the output and auxiliary nodes ($k_{23}$) to enable adaptation. Meanwhile, the lower ruggedness circuit has a more stable indirect buffer motif thanks to its interaction between the input and the auxiliary nodes ($k_{12}$) that distributes the adaptation mechanism across $k_{12}$ and $k_{23}$. This also reverses the direction of the signal response, as the signal RNA node 1 first binds with RNA node 2 and briefly frees up the output RNA node 3.

#### 3 Note 3: Motif clusters differing by ruggedness

Some of the motif groups are quite similar. For example, the stable motifs 2 and 3 differ in the strength of their node 2 self-interaction $k_{22}$, which is stronger than the $k_{33}$ interaction in motif 2 but similar or weaker in motif 3. Otherwise, both utilise self-interactions on all nodes to enact the adaptable mechanism, along with a $k_{13}$ interaction (as we have seen in Figure 2). The self-interaction on node 2 is essentially strengthening the stability of the adaptation mechanism enacted between nodes 1 and 3 by keeping node 2 out of the way from interfering. In comparison, $k_{22}$ is lacking in motif 5 and subsequently has a mix of stable and evolvable circuits. Motif 4 lacks yet another self-interaction by substituting the $k_{11}$ interaction with a $k_{23}$ interaction, meaning that node 2 acts to stabilise the output node 3 directly. The $k_{13}$ interaction in this motif cluster also only exists in quite a narrow range of bonding energies, which may explain why a substantial proportion of circuits from this cluster display high ruggedness. Finally, motif 1 takes this even further and has no self-interactions, enacting adaptation through $k_{12}$ and $k_{13}$ being weaker than $k_{23}$. This is the most unstable motif and can be rendered non-functional through perturbations in any interactions, while more stable motifs like 2 and 3 can afford mutations to some of their interactions (e.g. $k_{22}$) without losing adaptation.

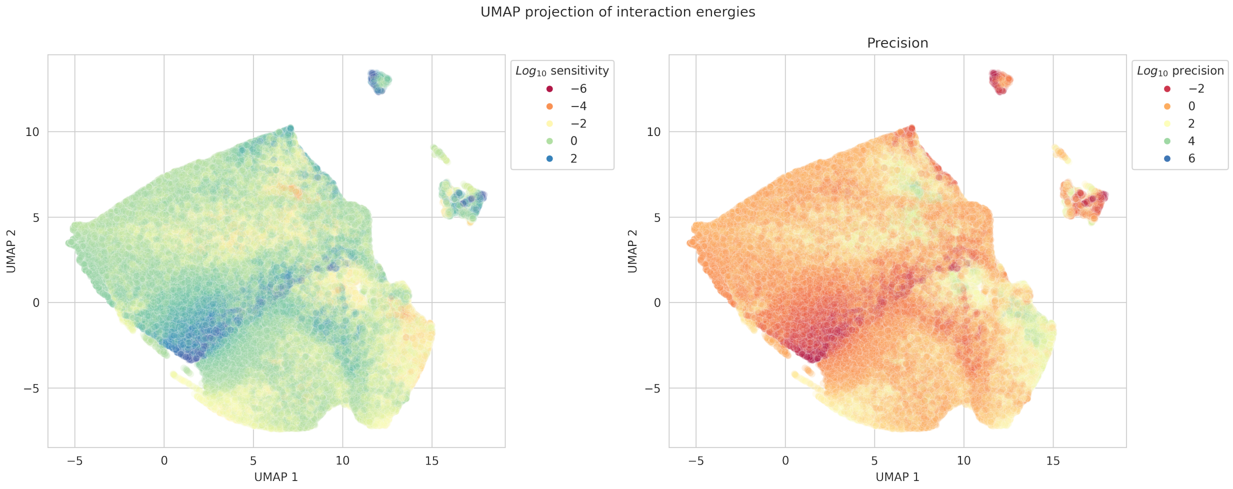


Supplementary Figure 1: UMAP dimensionality reduction of the simulated RNA circuits (trained directly on circuit topology) produces a latent space that reveals few features. Multiple regions correspond to high or low circuit functionality for both sensitivity and precision. While some trends do emerge just from features in the data, these are structures and rather grouped into one major cluster, with 2-3 very small satellite clusters. This warrants the use of a more sophisticated model like the CVAE to produce improved embeddings.

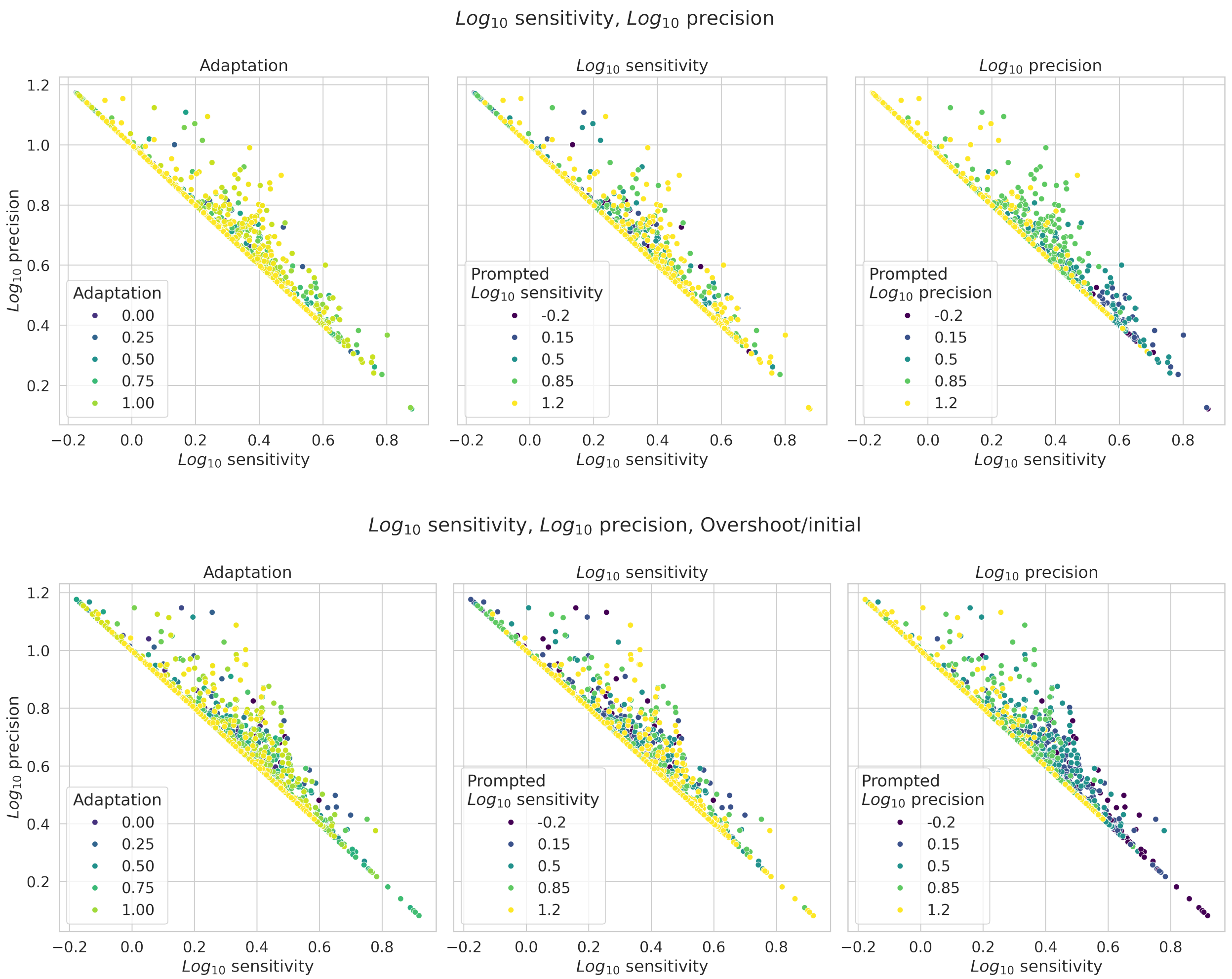


Supplementary Figure 2: For testing different objectives, some of the prompt distributions in Figure 3 were in the wrong order compared to the prompt that generated them. In the sensitivity vs. precision plots above, the prompt colours the samples, with the $log_{10}$ sensitivity and $log_{10}$ precision prompts being taken directly from the prompt and the adaptation prompt having been calculated from those two. For the top row, the model was trained with the objective [$log_{10}$ sensitivity, $log_{10}$ precision] and shows poor adherence to the adaptation and sensitivity prompts, but lines up much better with precision. The same can be said for the bottom row, where the model was trained with the objective [$log_{10}$ sensitivity, $log_{10}$ precision, overshoot/initial], though this one corresponds noticeably better to the sensitivity prompt. The precision portion of the prompt is still dominating the overall prompt. This could be because precision is achieved more simply compared to sensitivity and because sensitivity is the inverse of precision when there is no overshoot.

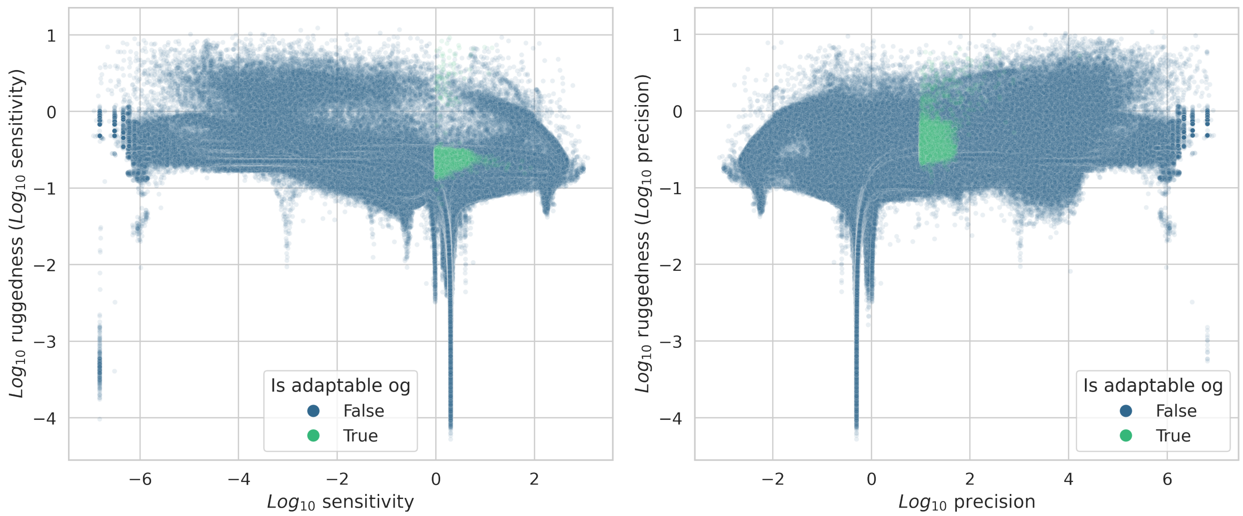


Supplementary Figure 3: The ruggedness of sensitivity and precision as the objective functions offer a comparison to looking at the ruggedess of adaptation compared to actual adaptation. Both sensitivity and precision plotted against their respective ruggedness show a mostly flat correlation with ruggedness, with some exceptions occurring for circuits with particularly low ruggedness near sensitivity $= 1$ and precision $= 1$. Ruggedness also slightly increases for some circuits with very low sensitivity and high precision. The adaptable region is shown in green as the original adaptable region, meaning the binary threshold definition of sensitivity $> 10$ and precision $> 1$ as opposed to a threshold in the custom adaptation function.

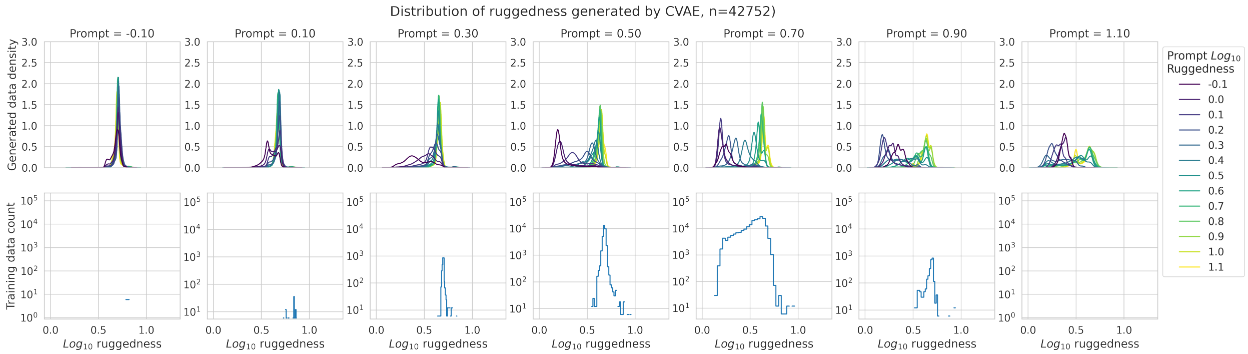


Supplementary Figure 4: The training data ruggedness is not evenly distributed across all levels of adaptation. In Figure 4, we showed the ruggedness prompt distributions for just the highest adaptation prompt. Here, we should the conditional distributions across 7 increasing adaptation ranges, with each distribution representing the true ruggedness of circuits generated for a ruggedness prompt. The adaptation prompt 0.70 is also the adaptation level around which training data is most plentiful.

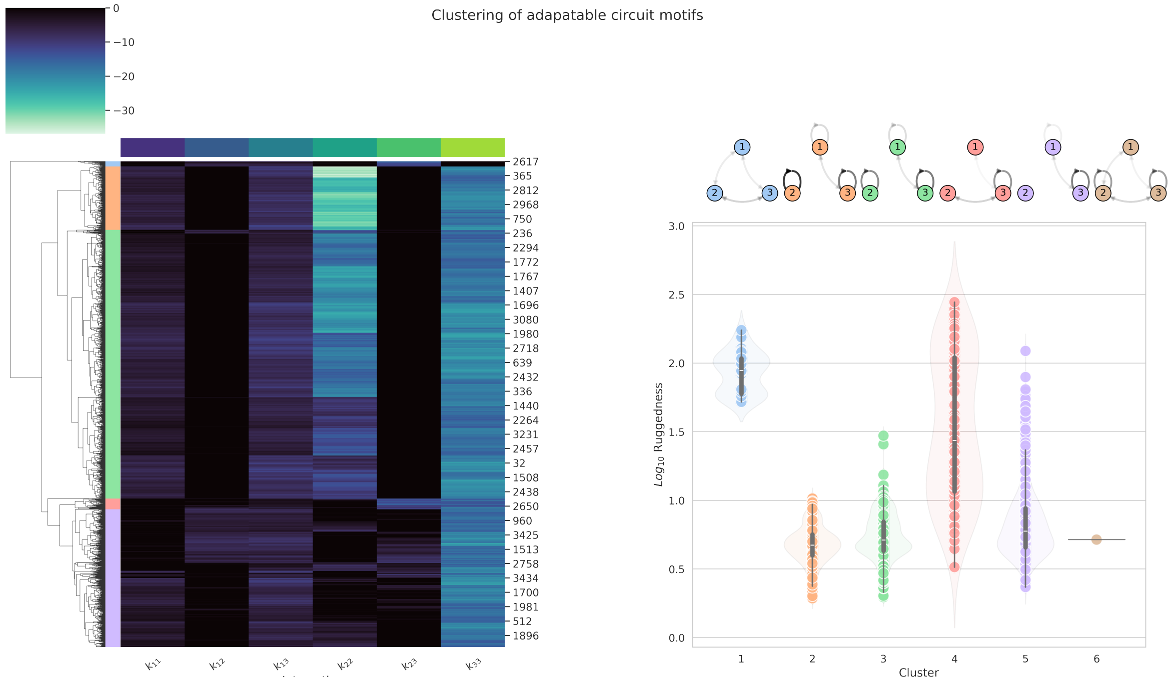


Supplementary Figure 5: Hierarchical clustering of the interactions of adaptable circuits generated by the CVAE (trained with adaptation and evolutionary ruggedness) identifies 6 motif groups. One of these (”6”) is only made up of one circuit, so we dropped this circuit in subsequent analyses.

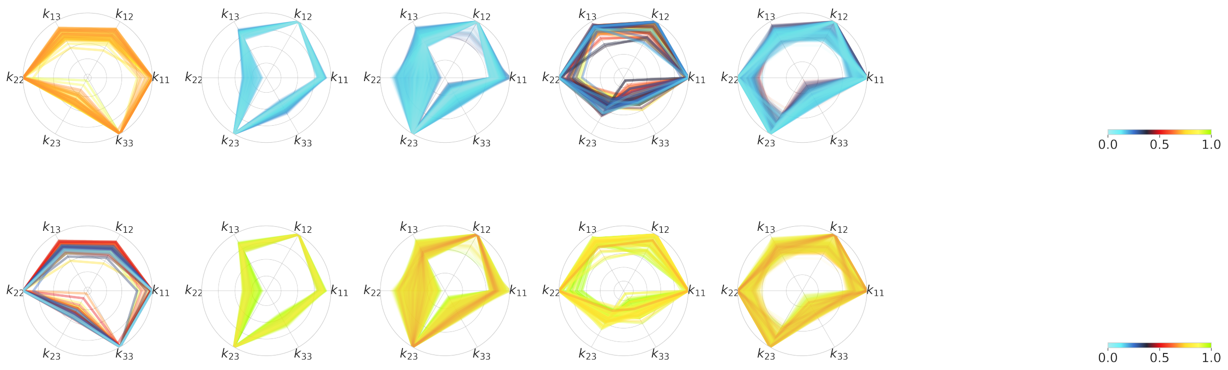


Supplementary Figure 6: Radar plots of the 5 adaptable motif groups (identified in Figure 4 and Supplementary Figure S5) with different evolutionary ruggedness show the connectivity between different interaction strengths and how these circuit constellations correspond to evolvability. The outside of each radar circle represents a binding interaction (minimum free energy) of 0, while the centre of the circle represents the maximum (approximately 35 kcal/mol), though we are only aiminig to show relative differences here. In the top row, circuit motifs are coloured by the $log_{10}$ ruggedness, normalised to the range [0, 1]. The bottom row shows the same motifs coloured by the ruggedness prompt that generated the circuit. Because of the normalisation, the exact values do not line up in terms of hue, but we are only interested in whether the CVAE identifies motifs that are more on the borderline of being evolutionarily stable. In the first motif for example, the circuits closer to the inside of the circle are more rugged, which is in line with the relative prompts of the CVAE. However, the CVAE also overestimated the region of stability for this motif. Something similar can be said for the fourth and fifth motifs, which also have less stable circuits on the inner edges of all motifs that are reflected by the prompt. Overall, the CVAE over-estimates the relative evolutionary stability of circuits, while prompts that are intended to be rugged also result in more rugged circuits.

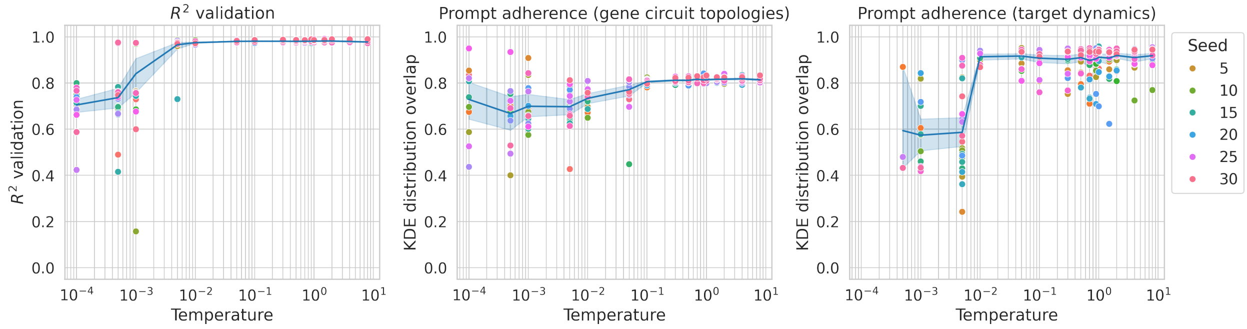


Supplementary Figure 7: Assessing contrastive loss: the temperature setting controls how strongly contrastive loss is applied, with a lower temperature punishing incorrect similarities between samples more. The training accuracy (measured by $R^2$) decreases with temperatures $< 10^{-2}$, so this is the minimum bound. Prompt adherence is assessed by how distinct the distributions of circuits are for our prompts, which is measured by the average overlap between a KDE prompt distributions and all other prompt distributions, averaged over all prompts. The distributions can be in terms of the circuit topologies, so how similar each unique interaction is distributed for each prompt, and in terms of the circuit's function, which has to be simulated once circuits are generated. While there is mild improvement in prompt adherence based on circuit topologies (e.g. a lower average KDE distribution overlap), the same is not true for the actual simulated circuit function of adaptation. In fact, higher temperature (weaker contrastive loss) actually results in some models with decreasing KDE distribution overlaps for adaptation, meaning that these models produce more distinct distributions for each prompt.
